# Energy inefficiency underpinning brain state dysregulation in individuals with major depressive disorder

**DOI:** 10.1101/2025.03.11.642725

**Authors:** Qianhui Liu, Hui Xiong, Weiyang Shi, Shiqi Di, Xinle Cheng, Nianyi Liu, Na Luo, Yu Zhang, Tianzi Jiang

**Author notes:** These authors contributed equally to the work (Qianhui Liu, Hui Xiong).

## Abstract

Disruptions in brain state dynamics are a hallmark of major depressive disorder (MDD), yet their underlying mechanisms remain unclear. This study, building on network control theory, revealed that decreased state stability and increased state-switching frequency in MDD are driven by elevated energy costs and reduced control stability, indicating energy inefficiency. Key brain regions, including the left dorsolateral prefrontal cortex, exhibited impaired energy regulation capacity, and these region-specific energy patterns were correlated with depressive symptom severity. Neurotransmitter and gene expression association analyses linked these energy deficits to intrinsic biological factors, notably the 5-HT2a receptor and excitatory-inhibitory balance. These findings shed light on the energetic mechanism underlying brain state dysregulation in MDD and its associated biological underpinnings, highlighting brain energy dynamics as a potential biomarker by which to explore therapeutic targets and advance precise interventions for restoring healthy brain dynamics in depression.

## Introduction

Major depressive disorder (MDD) is a prevalent mental illness associated with an increased risk of suicide. It is projected to become the leading global disease burden by 2030 [1–3]. Despite its significant impact, the underlying pathophysiology of MDD is not yet fully understood, although decades of neuroimaging studies have contributed valuable insights [4–9].

A major key for understanding MDD lies in how brain dynamics are altered in this disorder [10, 11]. It is well-established that the human brain is a highly complex and dynamic system [12], constantly exhibiting rich time-varying spatiotemporal patterns of brain states that represent recurrent configurations of neural activity and interactions, even at rest [13–16]. Normal brain functioning relies on the brain’s ability to flexibly transition between these states, supporting cognitive and affective processes [17–21]. In contrast, individuals with MDD seem to have difficulty regulating brain states effectively [22]. Specifically, empirical studies have indicated an impaired ability to maintain stable activity patterns, switch between states, and transition through them appropriately [23–26]. Moreover, disruptions in the dynamical reconfigurations of brain states have been observed in MDD, particularly in neural circuits related to sensory and motor functions, emotional processing, and cognitive control as well as in the default mode network [24–28]. These alternations to normal brain state dynamics were linked to symptom severity [23, 29, 30], rumination [31], and suicidality [32] in MDD. While these findings shed valuable insights into the pathophysiology of MDD, the underlying mechanisms that give rise to brain state dysregulations remain unclear, posing challenges to the development of therapeutic strategies aimed at effectively modulating brain state dynamics [10, 11, 33–35].

Since brain dynamics are fundamentally shaped by the underlying structural architecture, a promising approach for investigating these is network control theory (NCT) [10, 11, 36], which emphasizes the critical link between brain structure and function [13, 37–41]. Building on this, NCT can be used to model brain dynamics by integrating brain structural networks with functional activity patterns, to elucidate how network dynamics evolve through state space [42, 43]. This framework can be used to assess the average and modal controllability of specific brain regions or networks, which quantify the general ability of a certain brain region to influence other regions in a variety of ways [20]. It also allows researchers to profile the brain’s energy landscape by quantifying the energetic cost, or control energy, needed for transitions between recurrent brain states, enabling them to identify the states that are easier or harder to access and which brain regions need to be perturbed and to what extent to facilitate or maintain these transitions efficiently [21, 44–48].

While NCT has provided insights into the energy (in-)efficiency of state transitions in a growing body of neurological and psychiatric conditions, including epilepsy [44, 49], psychiatric disorders [50, 51], psychedelic effects [52], and neurodevelopment [53], its application in MDD remains inadequately explored. Previous studies applying NCT to MDD have primarily focused on structural or functional brain controllability to characterize the general ability of specific brain regions or networks to drive all possible states, thereby serving as potential biomarkers [54–59], and to search for potential stimulation targets for therapeutic interventions [60, 61]. However, these are not sufficient to answer questions about the control of specific states. Examinations of control energy in MDD through the framework of NCT could offer a means to uncover the dynamic mechanisms underlying the MDD brain state dysregulations. It is possible that state transition energies in MDD increase. Nevertheless, the energy landscape of brain activity states in MDD and how it might be impaired are still poorly understood. Moreover, whether NCT can offer a mechanistic explanation of why brain state dysregulations occur in MDD or which specific brain regions contribute to these dysregulations remains elusive.

The relationship between energy alterations and neurotransmitter systems and how these alterations interact with genetic factors are crucial for building a more biologically and mechanistically informed understanding of the pathophysiology of MDD. Neurotransmitter and genetic changes have long been recognized as key contributors to the biological underpinnings of MDD [4, 62–64]. Numerous studies bridging molecular and systems scales have shown that the imaging abnormalities observed in MDD are associated with specific neurotransmitter systems and gene expression profiles [65–70]. Recent advances in NCT have uncovered the neurophysiological correlates between control energy and both metabolic processes [49] and neurotransmitter perturbations [45, 52]. Building on these findings, we postulated that the neurotransmitters and genetic profiles might contribute mechanistic insights into control energy and, thereby, into the neurobiological underpinnings of MDD. At the time of our research this was yet to be determined.

To investigate these issues, a total of 890 participants identified with MDD and an equal number of matched healthy controls from the UK Biobank cohort [71–75] were included in this study (see Methods: Participants, and Supplementary Methods: Participants selection process). We began by identifying recurrent patterns of whole brain resting-state activity and examining how they differed between individuals with MDD and healthy controls (Fig. 1c). Our analysis revealed significant alterations in brain state dynamics in MDD. These were characterized by reduced state stability and increased switching rate. Then, we applied NCT to simulate the state transitions and estimate the associated energy costs (Figs. 1a and c), thereby probing the brain’s energy landscape and providing energetic evidence for the brain state disruptions observed in MDD. Our results revealed increased energy cost and reduced stability in MDD and demonstrated the association between altered energetic costs and changes in brain state transitions. We further explored these control profiles to identify the specific brain regions responsible for energy dysregulation (Figs. 1b and c), finding those that were critical for cognitive and emotional functioning and their association with depressive symptom severity. Finally, we investigated the relationship between these energy alterations and neurotransmitter engagement as well as with gene expression (Fig. 1c) to elucidate the biological mechanisms that contribute to the observed energy inefficiencies in MDD.

**Fig. 1.**
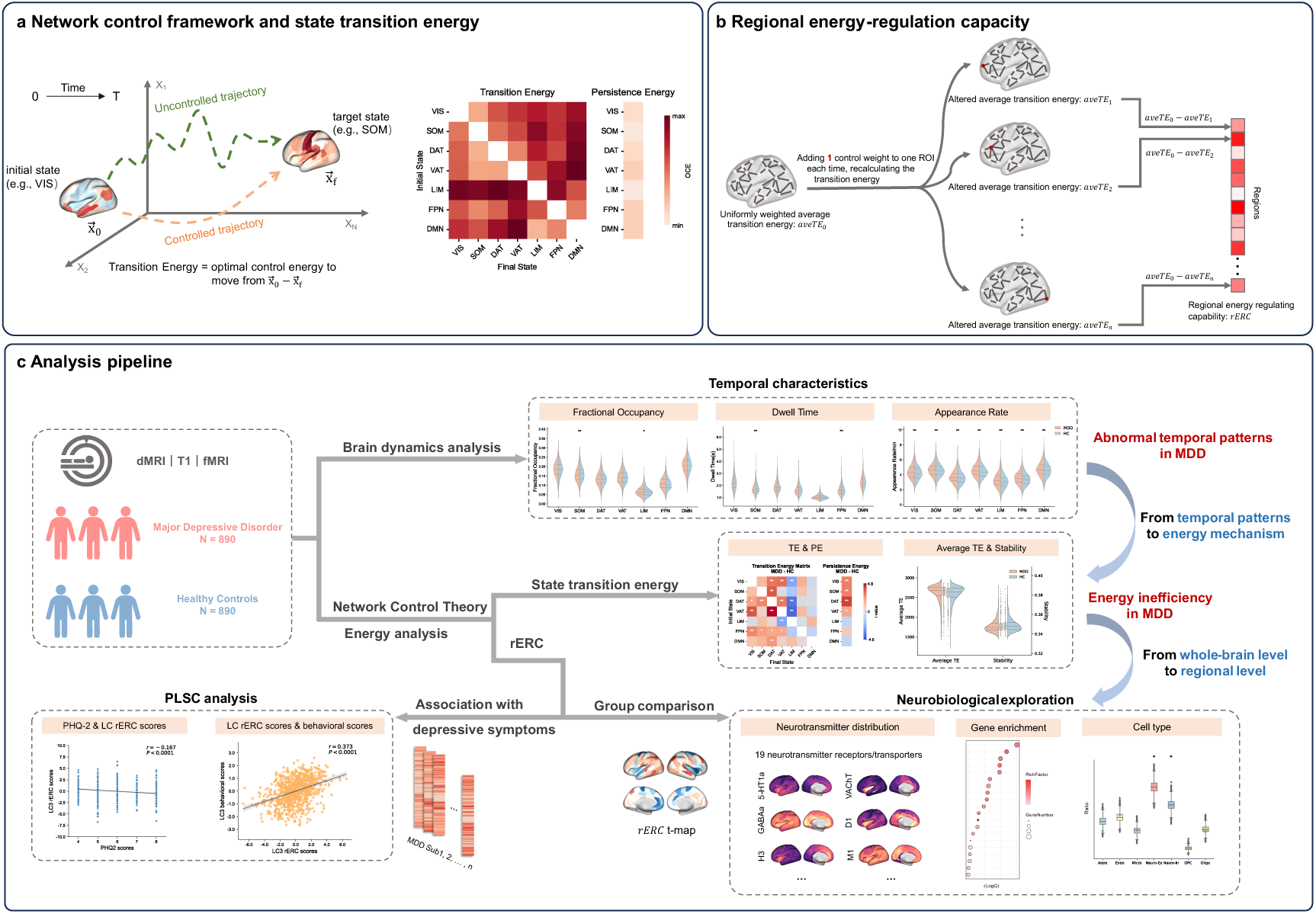
Schematic of methods and the study design. (a) A summary of how we used network control theory (NCT) to model brain state transitions and quantify the associated optimal control cost, as well as (b) to quantify the capacity of a certain region in regulating energy (namely regional energy regulation capacity, rERC) by changing its control input weights. (c) Schematic summary of the study design. Building on NCT, we investigated the energetic mechanism underlying disrupted brain state dynamics in MDD as well as its link to clinical, neurotransmitter, and genetic profiles. MDD, major depressive disorder; HC, healthy control; VIS, visual network; SOM, somatomotor network; DAT, dorsal attention network; VAT, ventral attention network; LIM, limbic network; FPN, frontoparietal network; DMN, default mode network; PE, persistence energy; TE, transition energy; dMRI, diffusion magnetic resonance imaging; fMRI, functional magnetic resonance imaging; PLSC, partial least squares correlation.

## Results

### Altered temporal patterns of brain states in MDD

The first step was to identify recurrent patterns of brain activity or states. In this study, we utilized a data-driven non-binary approach to define the brain states. This method preserves the intelligibility of intrinsic networks as states and is tailored to individual brain activity patterns [48]. Using this method with the Brainnetome atlas [76], seven centroid-like brain states that represent the activation of different resting-state intrinsic networks [77], namely the visual (VIS), somatomotor (SOM), dorsal attention (DAT), ventral attention (VAT), limbic (LIM), frontoparietal (FPN), and default mode (DMN) networks, were extracted from the resting-state fMRI BOLD activities (Fig. 2a, See Methods: Extraction of brain states).

**Fig. 2.**
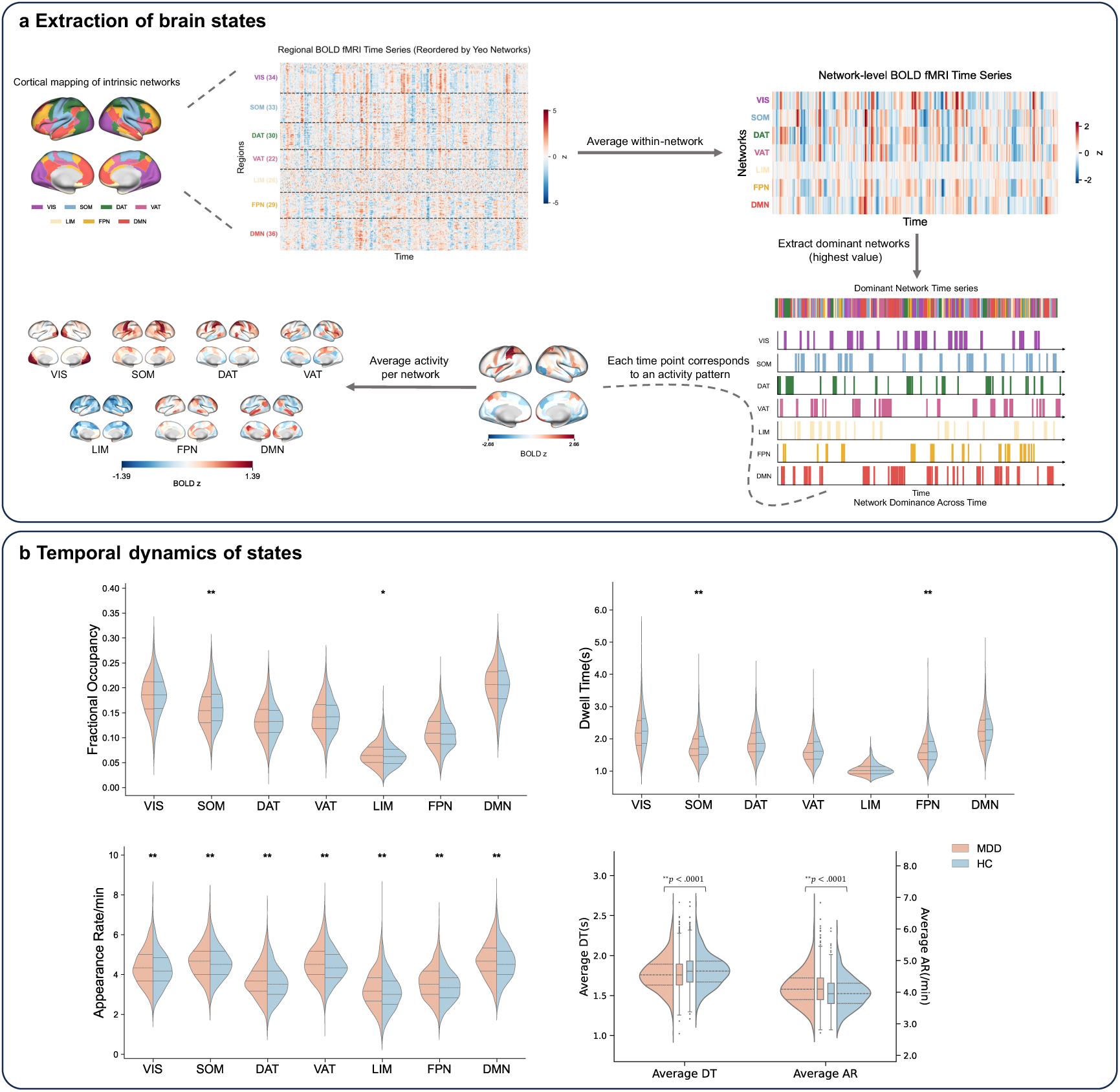
Altered temporal brain dynamics in MDD. (a) Schematic summary identifying recurrent states of brain activity, based on fMRI BOLD time series. (b) Comparison of temporal brain dynamics between MDD and HCs, including (top left) fractional occupancy, (top right) dwell time, and (bottom left) appearance rate. (bottom right) Average dwell time and average appearance rate across seven states were also compared. *uncorrected *p* < .05, ** *p* < .05 after Benjamini-Hochberg correction. MDD major depressive disorder; HC, healthy control; VIS, visual network; SOM, somatomotor network; DAT, dorsal attention network; VAT, ventral attention network; LIM, limbic network; FPN, frontoparietal network; DMN, default mode network; fMRI, functional magnetic resonance imaging; DT, dwell time; AR, appearance rate.

After identifying seven brain states for each subject, we were interested in whether and how brain state dynamics in the MDD patients differed from those of the healthy controls (HCs). Brain dynamics relies not only on the occurrence of various states but also on the capacity to transition flexibly between them [49]. Therefore, to systematically characterize the temporal sequence of brain states for each subject, we employed four related metrics: (1) fractional occupancy (FO), the proportion of the occurrences of a state (Fig. 2b, top left), (2) dwell time (DT), the mean duration (in seconds) that a given state was maintained (Fig. 2b, top right), (3) appearance rate (AR), how often each state appeared per minute (Fig. 2b, bottom left), and (4) transition probability, the probability of a given state switching to each of the other states (Fig. 3a).

**Fig. 3.**
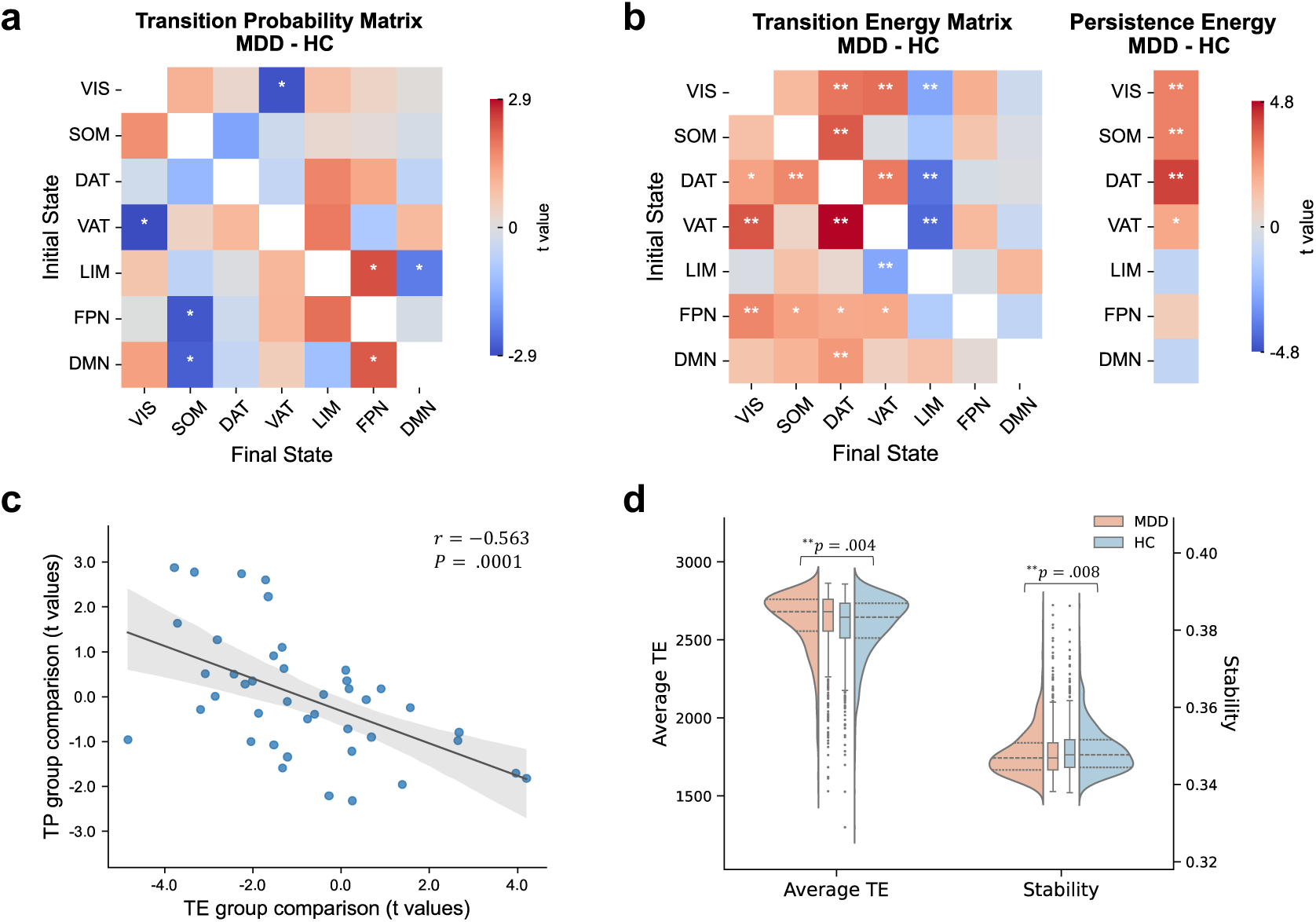
Energy inefficiency in MDD. (a) Case-control comparison of the empirically observed transition probabilities of different state pairs. (b) Case-control comparison of the transition and persistence energies of different state pairs using uniformly-weighted whole-brain inputs. Warm color means that the compared metric in MDD is greater than that in HC, and cold color on the contrary. (c) Between-group transition energy *t* values were significantly negative correlated with transition probability *t* values *r* = −0.563, *p* = .0001. (d) Case-control comparison of average transition energies across different states pairs and global control stability. *uncorrected *p* < .05, ** *p* < .05 after Benjamini-Hochberg correction. MDD major depressive disorder; HC, healthy control; VIS, visual network; SOM, somatomotor network; DAT, dorsal attention network; VAT, ventral attention network; LIM, limbic network; FPN, frontoparietal network; DMN, default mode network; TP, transition probability; TE, transition energy.

We found that for both the MDD patients and HCs, the brain most frequently occupies the state that represents the DMN (highest FO among the seven states), consistent with previous findings that highlight the DMN’s prominence during the resting state [78–80]. Notably, compared with the HCs, the individuals with MDD showed a significantly decreased FO in the SOM state (*t* = −2.958, *p*_FDR_ = .022) but a higher tendency of the FO in the LIM state (*t* = 2.354, *p*_FDR_= .065). Significant differences in DT were observed in both the SOM (*t* = −3.781, *p*_FDR_ = .001) and FPN states (*t* = −2.559, *p*_FDR_ = .037 ), with shorter durations in the MDD patients than in the HCs, indicating decreased stability of these two states in MDD. Additionally, the MDD patients demonstrated significantly higher AR across all states, revealing that states switch more often in MDD (VIS, *t* = 3.775, *p*_FDR_ = .0004; SOM, *t* = 2.650, *p*_FDR_ = .008; DAT, *t* = 2.755, *p*_FDR_ = .007; VAT, *t* = 3.345, *p*_FDR_ = .001; LIM, *t* = 4.114, *p*_FDR_ = .0001; FPN, *t* = 4.713, *p*_FDR_ < .0001; DMN, *t* = 3.383, *p*_FDR_ = .001). In line with the definitions of these metrics, the relationship between FO, DT, and AR can be understood as showing that a state with a high FO can result from either frequent transitions (high AR) or longer maintenance of that state (high DT) or a combination of the two. Thus, for the MDD patients, the reduced occupancy in the SOM state appears to be primarily due to its decreased stability, as reflected by its shorter DT, while the increased occupancy of the LIM state may have resulted from its increased rate of appearance (Fig. 2b). Furthermore, we averaged the DT and AR across the seven states and found that the MDD patients exhibited a shorter average DT (*t* = −3.888, *p*_FDR_ < .0001) and a higher average AR (*t* = 6.144, *p*_FDR_ < .0001) (Fig. 2b, bottom right). These findings suggest a different engagement of these intrinsic networks in the MDD patients than in the HCs during resting-state activity. Compared with a healthy system, the brains of the MDD patients may be less stable and more chaotic, as characterized by diminished state maintenance and more frequent state changes.

To assess the state transitions, we also calculated the transition probabilities for each individual and each condition and found that states switching to the FPN state were generally more probable in the MDD patients, while the HCs showed higher probabilities of transitioning to the SOM state (Fig. 3a). This is consistent with the earlier results shown in Fig. 2b, in which the HCs exhibited significantly increased FO in the SOM state whereas the MDD patients showed increased FO in the FPN state, which might have resulted from a higher AR. Although there were no significant group differences after multiple corrections, these results provide valuable trending insights into the altered engagement of key brain networks in MDD.

### Energy inefficiency in MDD: increased energy demands and reduced stability

Given that the brain state dynamics were significantly different between the MDD patients and the HCs, our next research question focused on the dynamical mechanisms underlying these state transitions, which we studied by quantifying the energetic effort associated with state transitions in the brain’s dynamic repertoire. Specifically, we aimed to investigate how individuals with MDD might exhibit energetic alterations during different state transitions and, thus, provide an energetic basis for the atypical temporal patterns of brain states observed in MDD.

To this end, we employed network control theory (NCT), which provides a framework for quantifying the ease of state transitions within a dynamical system [10, 11, 36, 42, 43]. Using NCT, we modeled the energy required for specific brain state transitions while considering the constraints of the underlying structural connectome (See Methods: Control energy and energy regulation capacity). For biological interpretability, a controlled path from an initial state to the final target state was designed to follow an optimal trajectory that balances the energy needed for the transition with the distance to the desired target state [21]. In this context, transition energy is defined as the optimal control energy necessary to facilitate transitions between various brain states. In the following analysis, the control energy required to remain in a given state is termed “persistence energy,” while “transition energy” specifically refers to the control energy associated with transitions between different states.

For each participant, we first calculated the optimal control energy required to transition between each pair of brain states by applying uniform control input weights across all brain regions. Transitions are significantly less energetically costly in the structural connectome than in their null models (See Supplementary Methods: Network null models), suggesting that structural connectome wiring supports effective transitions between brain states (Fig. S4). We then performed a Spearman correlation analysis between transition energy and transition probability, revealing a strong negative correlation in both the MDD (*r* = −0.68, *p* < .0001; Fig. S5) and HC groups (*r* = −0.70, *p* < .0001; Fig. S5). This finding aligns with the idea that the brain prefers trajectories through state space that require little input energy [18, 48], supporting the use of transition energy as an effective indicator of transition ease.

In comparing the transition energies between the two groups, we found that transition generally demanded more energy in the MDD patients than in the HCs for most state pairs (Fig. 3b), suggesting decremented efficiency in the diseased brain [49, 51]. Interestingly, transitioning into the LIM state required less energy in the MDD patients, which provides an energetic explanation for the observed changes in temporal patterns of brain dynamics. Specifically, the MDD patients exhibited a higher occupancy and appearance rates for the LIM state (Fig. 2b) and a greater likelihood of switching to this state (Fig. 3a) than the HCs, likely due to the lower energetic cost associated with such transitions. In contrast, the HCs tend to experience more frequent occurrences and appearances of the SOM state, as these transitions are less energy demanding (Figs. 2b and 3b). Furthermore, we found that the group differences in transition energies were significantly negative correlated with their transition probability differences (*r* = −0.563, *p* = .0001, Fig. 3c). These findings indicated that differences in energetic costs can explain how the brain states shifted in the MDD patients, when compared with the HCs.

To further explore system dynamics, the transition energies across different state pairs were averaged, and MDD exhibited a higher mean energetic cost to facilitate these state switches (*t* = 2.885, *p* = .004 ; Fig. 3d, left). This implies increased barriers in the energy landscape in MDD. We also introduced a global stability metric, and found that MDD showed significantly reduced control stability (*t* = −2.638, *p* = .008; Fig. 3d, right). This finding corroborates our previous inferences from the temporal dynamics of MDD in Fig. 2b. Together, these results indicate energy inefficiency in MDD.

### Impaired regional energy regulation capacity in MDD associated with symptom severity

Previously, we observed an elevated energy burden in the MDD patients compared with the HCs. Next, we aimed to explore whether this phenomenon could be attributed to an aberrant energy regulation of specific brain regions.

One crucial component of control dynamics is the control set, namely, the set of control nodes or brain regions where control inputs are injected to induce transitions between brain states [43]. In these initial analyses, we used a uniform full control set, treating all brain regions equally in terms of their influence on system dynamics. In the ones that follow, we utilized region-specific heterogeneous control weights [45, 52] to examine the regulatory roles of specific brain regions in controlling brain dynamics. Specifically, we increased the control weight of a selected brain region by one while keeping the control weights of all the other regions unchanged (Fig. 1b; see Methods for details). Since increasing the total quantity of the control inputs necessarily reduces energy [42], this adjustment of the control weight allowed us to quantify the decrease in overall transition energy compared with the uniform control set. The degree of energy reduction attributed to a specific brain region reflects its capacity to regulate the energy cost within the system. We repeated this process for each brain region, generating a region-wise vector of regional energy regulation capacity (rERC) for each participant.

We then conducted a case-control comparison of the rERC between the MDD patients and the HCs, and the resulting statistical *t*-map is presented in Fig. 4a. Our analysis identified ten brain regions that displayed significant group differences, as reflected by the decreased rERC attributed to these regions in the MDD patients (Fig. 4b, Table 1). This suggests that the overall energy inefficiency observed in MDD may stem from the impaired energy regulation ability in these brain regions. Of these regions, three were classified within the DMN, involving one subregion of the left superior frontal gyrus, SFG_L_7_3 (A9l, lateral area 9), as well as two subregions of the cingulate gyrus, CG_L_7_3 (A32p, pregenual area 32) and CG_L_7_7 (A32sg, subgenual area 32). Notably, the SFG_L_7_3 (A9l, lateral area 9) is included in the left dorsolateral prefrontal cortex (dlPFC), which is an USFDA-approved stimulation target in transcranial magnetic stimulation treatment for depression [3, 81, 82]. Additionally, two subregions of the insular gyrus and three subregions within the SOM were identified (Table 1).

**Fig. 4.**
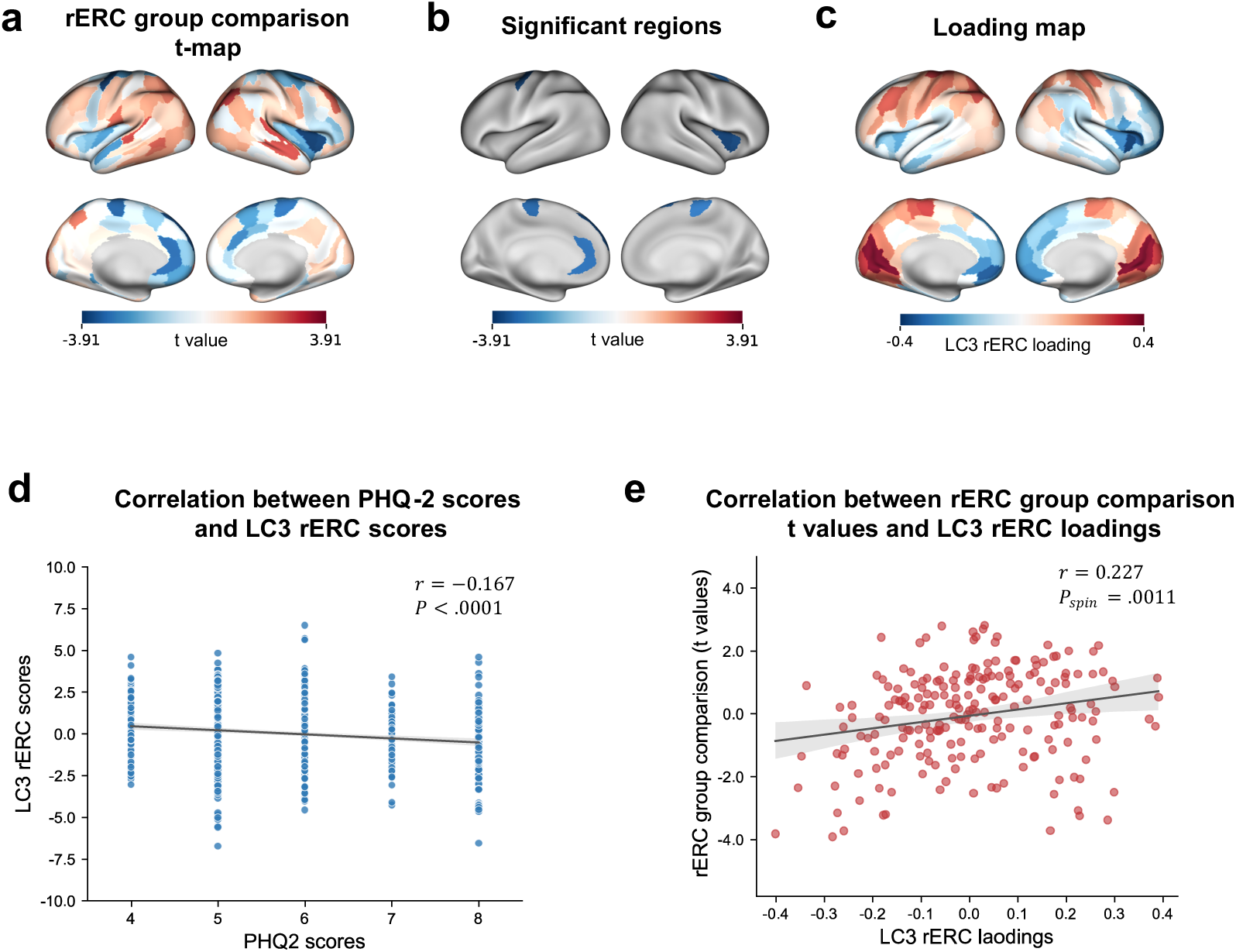
Impaired rERC in MDD and its link to clinical PHQ-2 scores. (a) Region-wise statistical comparisons of rERC between MDD and healthy controls. (b) Ten identified regions showing statistically significant group differences in rERC patterns (*p*_FDR_ <.05). To assess the relationship between rERC patterns in MDD and the severity of depressive symptoms (quantified by PHQ-2 scores), a PLS correlation analysis was applied. (c) The third PLS latent component (LC3) loadings of rERC were mapped to the cortical surface. (d) The LC3 rERC scores were significantly negatively correlated with the PHQ-2 scores (*r* = −0.167, *p* < .0001). (d) The between-group rERC *t* values were significantly positively correlated with the LC3 rERC loadings ( *r* = 0.227, *p_spin_* = .0011). MDD, major depressive disorder; rERC, regional energy regulation capacity; PLS, partial least squares; LC, latent component; spin, spin tests.

**Table 1.** Significant regions showing diminished rERC inc individuals with MDD compared with healthy controls (*p*_FDR_ < .05). VIS, visual network; SOM, somatomotor network; DAT, dorsal attention network; VAT, ventral attention network; LIM, limbic network; FPN, frontoparietal network; DMN, default mode network; FDR, false discovery rate.

| Gyrus | Label | Name | Anatomical and modified cyto-architectonic descriptions | Intrinsic network | *t*-stats | *p*<sub>FDR</sub> |
| --- | --- | --- | --- | --- | --- | --- |
| Superior frontal gyrus | 5 | SFG_L_7_3 | A9l, lateral area 9 | DMN | -3.7249 | .0112 |
|  | 8 | SFG_R_7_4 | A6dl, dorsolateral area 6 | DAT | -3.2229 | .0374 |
| Precentral gyrus | 55 | PrG_L_6_2 | A6cdl, caudal dorsolateral area 6 | DAT | -3.7099 | .0112 |
|  | 59 | PrG_L_6_4 | A4t, area 4(trunk region) | SOM | -3.2433 | .0374 |
| Paracentral lobule | 67 | PCL_L_2_2 | A4ll, area 4, (lower limb region) | SOM | -3.3776 | .0314 |
|  | 68 | PCL_R_2_2 |  | SOM | -3.0681 | .04593 |
| Insular gyrus | 168 | INS_R_6_3 | dIa, dorsal agranular insula | VAT | -3.8142 | .01122 |
|  | 174 | INS_R_6_6 | dId, dorsal dysgranular insula | VAT | -3.9075 | .01122 |
| Cingulate gyrus | 179 | CG_L_7_3 | A32p, pregenual area 32 | DMN | -3.1945 | .03742 |
|  | 187 | CG_L_7_7 | A32sg, subgenual area 32 | DMN | -3.1497 | .03877 |

We further examined the relationship between rERC patterns in the MDD patients and the severity of depressive symptoms (quantified by PHQ-2 scores) using partial least squares (PLS) correlation analysis [83], while controlling for age, sex, and assessment center as covariates (Fig. S7). The results revealed that the third PLS latent component (LC3) significantly explained 22.44% of the covariance (*p* < .0001), and higher LC3 rERC scores significantly correlated with lower PHQ-2 scores (*r* = −0.167, *p* < .0001; Fig. 4d; see Supplementary materials for more details). This finding underscores a potential link between improved energy regulation capacity and reduced depressive symptom severity. Further analysis showed a significant positive correlation between the rERC LC3 loadings (Fig. 4c) and the case-control rERC *t*-map ( *r* = 0.227, *p*_$%&’_ = .0011 ; Fig. 4e). This alignment indicates that the contribution of each brain region in predicting symptom severity is captured by the rERC alterations, highlighting rERC patterns as promising biomarkers for depression.

### Energy regulation deficits associated with brain-wide neurotransmitter distribution

Neurotransmitter alterations are implicated in the pathophysiology of MDD [62]. Therefore, we next sought to provide a biological interpretation of the deficits in energy regulation in MDD by systematically probing the relationship of this regulation with the brain’s underlying neurotransmitter engagement. Specifically, we investigated the spatial association between the neurotransmitter receptor/transporter density maps and the case-control rERC *t*-map (Fig. 5b). A total of 19 neurotransmitter receptors and transporters (Fig. 5a) were considered here, including dopamine, norepinephrine, serotonin, acetylcholine, glutamate, γ-aminobutyric acid (GABA), histamine, cannabinoid, and opioid, across nine different neurotransmitter systems [84]. Notably, our analysis revealed that of the neurotransmitter systems examined, the serotonin (5-HT) receptors, specifically 5-HT2a (*r* = 0.28, *p_spin_* < .0001, after Bonferroni correction), exhibited the most significantly positive correlation with the between-group differences in the rERC (Fig. 5b). These findings support the hypothesis that the neurotransmitter system serves as an intrinsic biological factor contributing to the observed energy dysregulation in MDD.

**Fig. 5.**
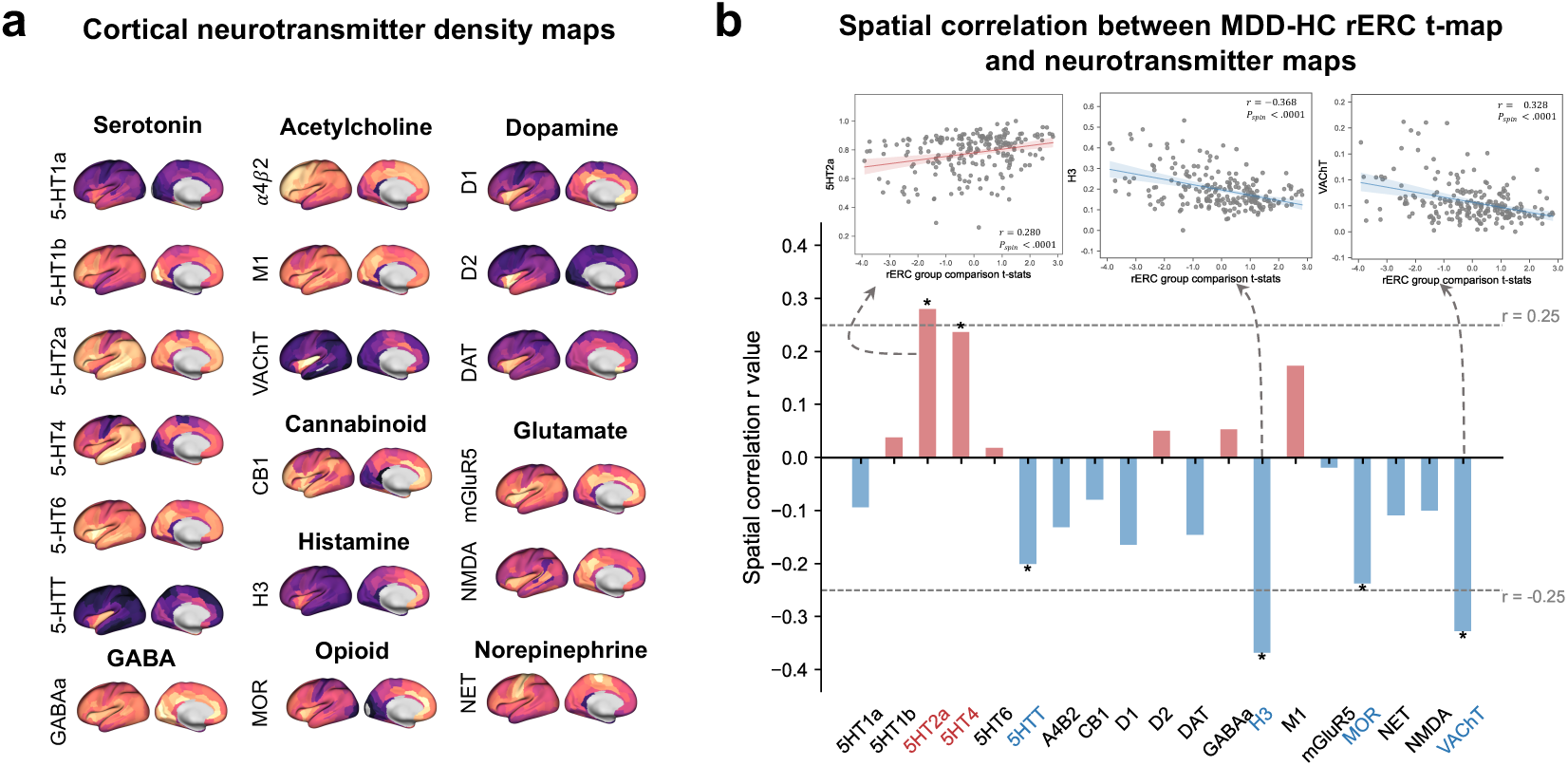
Association between brain-wide neurotransmitter distribution and energy regulation deficits in MDD. (a) The empirical spatial distribution of each receptor and transporter across nine neurotransmitter systems is mapped to the cortical surface using Brainnetome atlas. These include serotonin (5-HT_1a_, 5-HT_1b_, 5-HT_2a_, 5-HT4, 5-HT6, 5-HTT), GABA (GABA_a_), acetylcholine (α4β2, M1, VAChT), dopamine (D1, D2, DAT), cannabinoid (CB1), histamine (H3), glutamate (mGluR5, NMDA), opioid (MOR) and noradrenaline (NET). Raw PET data were obtained from [84]. (b) Distribution of the correlation coefficients between the rERC *t*-map and the spatial neurotransmitter maps. Relatively strong significant positive/negative correlation were found for 5-HT_2a_ (*r* = 0.28, *p_spin_* < .0001) / H3(*r* = −0.368, *p_spin_* < .0001) and VachT (*r* = −0.328, *p_spin_* < .0001). * *p* < .05 after Bonferroni correction. MDD, major depressive disorder; HC, healthy control; rERC, regional energy regulation capacity; spin, spin tests.

### Alterations in energy regulation ability associated with gene expression profiles

To further investigate the underlying biological mechanisms of the altered energy regulation patterns in MDD, we utilized PLS regression [68] to explore the spatial relationship between the rERC *t*-map and patterns of brain-wide gene expression from the Allen Human Brain Atlas (AHBA) [85, 86]. The findings revealed that the first PLS component (PLS1) explained 16.8% of the variance (*p_spin_* = .0042), and that the PLS1 gene expression scores were significantly spatially correlated with the case-control *t*-map (*r* = 0.328, *p_spin_* = .0001; Fig. 6a). This suggests that the largest regional variations in the transcriptional architecture of the human cortex can be effectively captured in the between-group rERC differences.

**Fig. 6.**
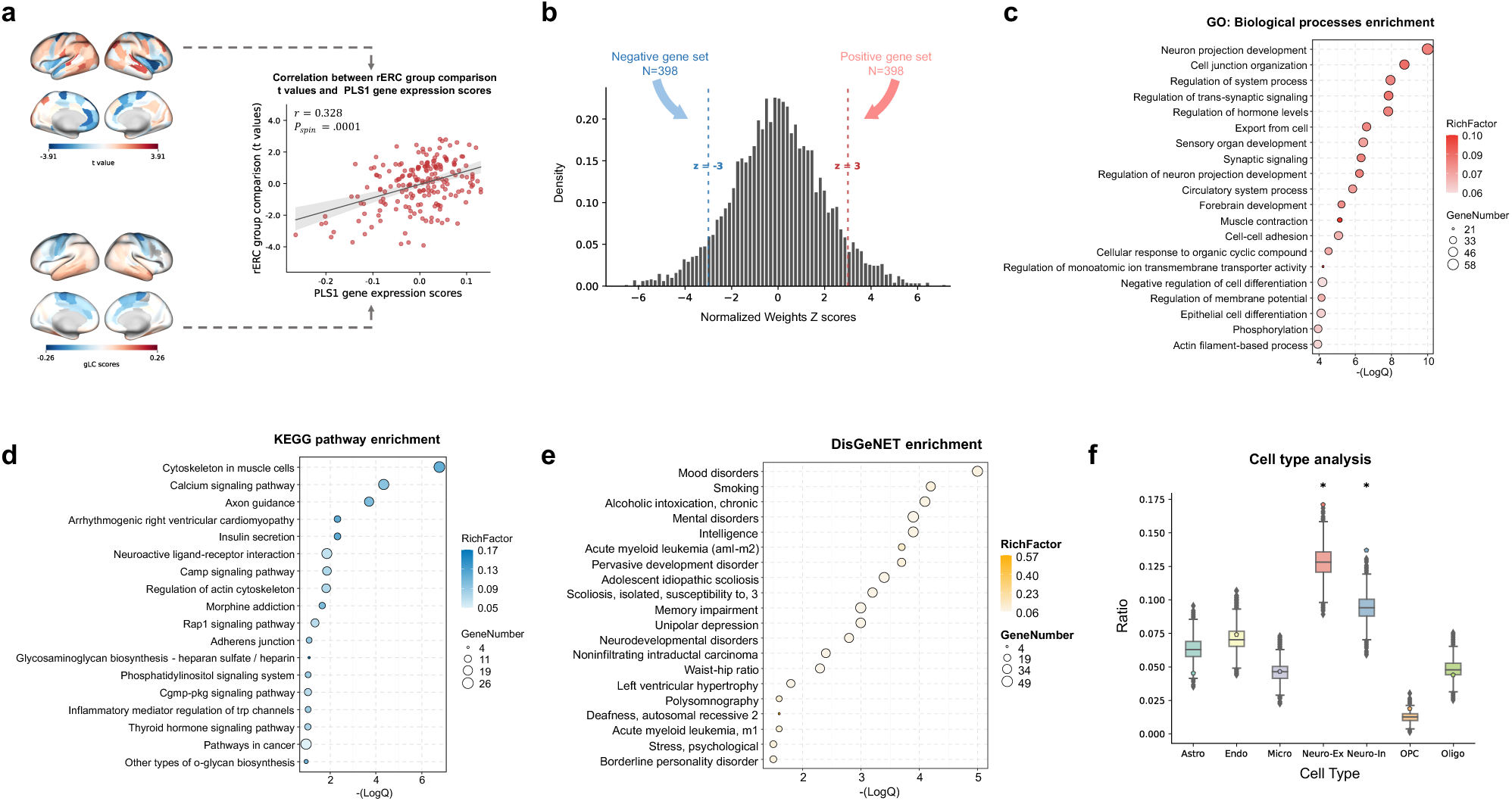
Gene expression profiles related to energy regulation deficits in MDD. (a) The PLS1 gene expression scores were significantly spatially correlated with the case-control rERC *t* values (*r* = 0.328, *p_spin_* = .0001). (b) Distribution of normalized PLS1 gene weights. 796 significant PLS1 genes (|Z| > 3) were selected for further enrichment analysis. Top 20 of the (c) Gene ontology (GO) Biological Processes functional annotations, (d) the KEGG pathway, and (e) DisGeNET enrichment. (f) Cell-type specific expression analysis of the PLS1 genes. * *p* < .05 after Bonferroni correction. MDD, major depressive disorder; rERC, regional energy regulation capacity; GO, gene ontology; KEGG, Kyoto Encyclopedia of Genes and Genomes; PLS, partial least squares; LC, latent component; Astro, astrocyte; Endo, endothelial; Micro, microglia; Neuro-Ex, excitatory neuron; Neuro-In, inhibitory neuron; Oligo, oligodendrocyte; OPC, oligodendrocyte precursor cell; spin, spin tests.

We then ranked the genes based on their normalized PLS1 weights (i.e., Z scores of PLS1 loadings), and a threshold of |Z| > 3 was applied (*p*_FDR_ < .05). The retained 796 genes were further analyzed using Metascape [87] for enrichment analysis (Fig. 6b). The gene ontology (GO) enrichment analysis of the biological processes identified terms related to cellular processes, such as neuron projection development and cell junction organization, as well as biological regulation, such as regulation of system process and regulation of trans-synaptic signaling (Fig. 6c). The KEGG Pathway analysis revealed significant enrichment in pathways associated with neural signal transduction, such as the calcium signaling pathway and cAMP signaling pathway, as well as pathways involved in the development and regeneration of organismal systems, such as axon guidance. Additionally, pathways associated with signaling molecules and interactions, such as neuroactive ligand-receptor interaction, were significantly enriched (Fig. 6d). The disease enrichment analysis using DisGeNET [88] demonstrated a strong association of the selected gene set with categories closely related to MDD, such as mental disorders and unipolar depression (Fig. 6e). Furthermore, a cell type deconvolution analysis [89] was performed to assess the preferential expression of these genes and showed that they are preferentially expressed in specific cell types with significant expression in excitatory (Neuro-Ex) (*p*_bonf_ = .0007) and inhibitory (Neuro-In) neurons (*p*_bonf_ = .0007) (Fig. 6f).

## Discussion

Understanding how brain state dynamics is altered in individuals with major depressive disorder (MDD) and comprehending the underlying mechanisms that alter the dynamics in MDD are essential for elucidating the pathophysiology of MDD and further developing effective therapeutic interventions.

This work investigated the energetic mechanism underlying disrupted brain state dynamics in MDD along with its associated biological underpinnings. Our findings revealed significant alterations in brain state dynamics in MDD, which were characterized by decreased state stability and increased state-switching frequency. Then, building on network control theory, modeling the brain state transitions revealed a more challenging energy landscape for individuals with MDD and provided mechanistic insights into the observed brain state disruptions. By analyzing the regional energy regulation capacity (rERC), we also identified key brain regions, including subregions of the left dorsolateral prefrontal cortex (dlPFC), subgenual cingulate cortex, and insular cortex, that contribute to the energy inefficiency in MDD and further showed the predictive value of rERC for assessing depressive symptom severity. Further analyses established associations between the regional energy profiles using both neurotransmitter and gene expression profiles and identified neurotransmitter systems, biological processes, signaling pathways, and cell types that are closely linked to the known pathophysiological mechanisms of MDD, providing further insight into the molecular basis for the observed energy regulation deficits in MDD.

Analyses of the brain state dynamics revealed significant changes in brain network activity in MDD, particularly within the SOM, FPN, and LIM networks, potentially identifying the disrupted brain functions associated with motor and cognitive control and with emotion regulation in depression [6, 28, 90]. This is consistent with existing studies reporting relevant alterations in these network dynamics [6, 24, 26, 28, 29] and seems to reflect the hallmark symptoms of MDD such as emotional dysregulation, fatigue or loss of energy, and psychomotor retardation [1]. The results also indicated that the brain in MDD functions as an unstable system with a lack of stability in maintaining brain states and an increased rate of state transitions. Modeling the brain state controls showed that this brain state dysregulation, a pathological trait of MDD, is shaped by the control costs. Moreover, MDD exhibited reduced stability and elevated energy barriers that impede the brain in navigating its dynamic landscape effectively, a finding which aligns with disease-related energy inefficiency [49–51]. Prior studies have linked control energy to oxygen and glucose metabolism, which the brain consumes to produce metabolic energy to support neural dynamics [48, 49]. The brain, as an economic system, balances costs with functional efficiency [91]. Thus, in MDD, it is possible that failing to meet these increased energy needs may destabilize the neural system, contributing to the wide-ranging cognitive and affective deficits observed in this disorder [90, 92–94].

From the broader perspective of controlling brain dynamics, identifying the brain regions that can be perturbed to facilitate state transitions in the most efficient way is critical for developing targeted treatments. To this end, we modeled region-specific influences in system dynamics to assess each region’s capacity for regulating energy (rERC). Our analysis identified brain regions, including the left dlPFC, subgenual cingulate, and insular cortex, all of which showed significantly reduced ability in regulating energy cost in MDD. These regions are known for their roles in cognitive and emotional processing [6, 28, 90]. Moreover, the region-specific energy patterns were predictive of depressive symptom severity, suggesting that the energy inefficiency observed in MDD may stem from impaired energy regulation in these key regions, and thereby contribute to the associated brain dysfunctions reflected in abnormal brain state dynamics. Additionally, structural and functional abnormalities in these regions, such as changes in gray matter volume, cortical thickness, and connectivity, have been widely reported [4, 9, 95–99]. Our results align with these observations and suggest that rERC patterns can well-capture these structural or functional aberrations. This further supports the potential for targeting these regions for stimulation-based interventions, such as transcranial magnetic stimulation (TMS) targeting the left dlPFC [3]. The subgenual cingulate cortex [33, 100, 101], anticorrelation between left dlPFC and subgenual cingulate [34, 102, 103], and insula-dlPFC connectivity [104, 105], have also been identified as successful therapeutic targets. These findings underscore the potential of energy-based patterns to provide promising biomarkers and target indicators for depression, offering insights into the mechanisms of stimulation-based treatments. By stimulating regions with impaired energy regulation capacity, external interventions may help restore the brain’s ability to manage control costs. Building on these, the use of healthy energy patterns as a reference could provide a framework for personalized assessments in MDD. Furthermore, linking brain states and their associated energy dynamics to mood states could offer mechanistic insights into mood-state-dependent stimulation, showing promise in precise neuromodulation tailored to specific mood states [9, 33].

Neurobiological interpretations of energy regulation deficits in MDD can be further explored through brain-wide neurotransmitter and gene expression association analyses. We found the strongest positive correlation between the between-group rERC patterns and the spatial distribution of the serotonin 5-HT2a receptor [84]. This suggests that reduced 5-HT2a receptor levels may contribute to impaired energy regulation in MDD, aligning with the serotonin hypothesis that a decrease in serotonin level is a risk factor for depression [62]. Notably, psychedelic treatments for depression, such as psilocybin, have shown promising antidepressant effects because of their agonist actions at the 5-HT2a receptors [3, 62]. A previous study demonstrated that serotonergic psychedelics, such as LSD and psilocybin, could flatten the brain’s control energy landscape [52]. These findings may imply that serotonergic psychedelics lower the increased energy barriers in MDD. Moreover, both theoretical and experimental studies have proposed that the brain’s general ability or readiness to transition between states is critically modulated by serotonergic signaling [10, 106]. In line with these evidences, our findings highlight the potential role of the serotonergic pathway in contributing to the energy dysregulation observed in MDD, supporting this pathway as critical to both MDD pathology and its treatment.

A neuroimaging-transcriptomic analysis revealed a significant spatial concordance between the changes in the rERC pattern and gene expression profiles across the cerebral cortex. A gene enrichment analysis identified several significant functional categories associated with MDD-related genes, including neuronal development, calcium signaling, and axon guidance. These findings are consistent with existing literature, indicating that neuronal impairment and maldevelopment may play a crucial role in the pathophysiology of MDD [67, 107, 108]. Disruptions in calcium signaling have also been implicated in the neural and behavioral maladaptation associated with MDD [109, 110]. Moreover, cell type enrichment revealed significant involvement of excitatory and inhibitory neurons, emphasizing the importance of maintaining the excitation-inhibition balance for proper brain function [65]. Dysregulation in this balance has been identified as a probable mechanism underlying depression [62]. These results indicate that the balance in neural activity and pathways is disrupted in MDD. A disease enrichment analysis revealed a strong association with mood disorders, further supporting the relevance of the molecular processes identified in this study for understanding the neurobiological mechanisms underlying depression.

While our findings offer valuable insights, several methodological considerations should be noted. First, our modeling of neural dynamics assumes linearity and time invariance, following previous studies [11, 20, 42, 43, 46, 49–52]. Although brain dynamics are inherently nonlinear, this can be justified because linear models provide reasonable approximations of brain dynamics [37, 111] and have been shown to outperform some nonlinear models in describing macroscopic brain activity [112]. In addition, a controllable linearized system can be locally controllable in nonlinear situations [113]. Nevertheless, as our understanding of the brain’s nonlinearity deepens, further work should explore the nonlinear dynamics for a more comprehensive understanding of the brain processes in MDD [114, 115]. Second, the structural connectivity used in our study was derived from diffusion tractography, which serves as an approximation of the brain’s true structural connectome underpinning functional brain dynamics. While this is the current state-of-the-art method, we are aware that it has an inherent limitation in fully capturing the real structural connectome of the brain [116]. Incorporating alternative connectivity representations may offer complementary perspectives [43, 49]. Third, the neurotransmitter and gene expression analyses were based on publicly available data rather than on participant-specific data. The results obtained are preliminary and unconclusive, and future studies should aim to integrate personalized data for more precise biological interpretations. Last, while our study benefitted from a large sample size, further clinical or longitudinal datasets are needed to validate these findings and establish their utility as biomarkers or therapeutic targets.

In summary, this study provided a comprehensive investigation into the energetic and biological mechanisms underlying altered brain state dynamics in MDD, highlighting the potential of brain energy dynamics as a source of biomarkers and as a framework for exploring therapeutic targets. Our findings offer a novel perspective on the neurobiological underpinnings of MDD and advance our ability to control its brain dynamics, paving the way for developing personalized, precise therapeutic strategies aimed at restoring healthy brain dynamics for individuals with depression.

## Methods

### Participants

This study utilized data from the UK Biobank (application number #88660) and included 38,979 participants with complete imaging data across T1-weighted MRI, rs-fMRI, and dMRI [71–75]. After excluding individuals with neurological or psychiatric conditions known to impact cognitive function, as well as those with incomplete data, a strict screening process was employed to identify 890 participants with major depressive disorder (MDD) based on their history of depression and recent core depressive symptoms. An equal number of healthy controls were matched to the MDD participants based on age, sex, and assessment center, ensuring a well-balanced comparison in this case control study (Table S1). Further details about the participant selection processes along with sample demographic and clinical characteristics are provided in the supplementary materials (see Supplementary Methods: Participants selection process).

### MRI acquisition and processing

The acquisition and preprocessing of MRI data were conducted by UK Biobank following their standardized imaging protocols [72, 74]. All imaging data were collected on Siemens Skyra 3T scanners running VD13A SP4, which were equipped with 32-channel RF receive head coils. T1-weighted structural images were acquired using a 3D MPRAGE sequence (orientation, sagittal; resolution, 1×1×1 mm; field of view (FOV), 208×256×256 matrix; repetition time (TR), 2,000 ms; echo time (TE), 2.01 ms) with a scan duration of 5 minutes. Resting-state functional MRI was performed using a gradient-echo EPI sequence (TR, 735 ms; TE, 39 ms; resolution, 2.4×2.4×2.4 mm; FOV, 88×88×64 matrix; flip angle, 52°) with a multislice acceleration factor of 8 and fat saturation, collecting 490 volumes over 6 minutes. Diffusion MRI data was acquired using a spin-echo EPI sequence (resolution, 2×2×2 mm; FOV, 104×104×72 matrix) with a multislice acceleration factor of 3. The diffusion protocol comprised 100 distinct diffusion-encoding directions (50 directions each at b = 1000 and 2000 s/mm²) and 8 b = 0 images (including 3 blip-reversed), with a total acquisition time of 7 minutes. MRI scans marked as corrupted or unusable by UK Biobank’s quality control procedures were excluded from further analyses.

Structural connectome and functional time series processing were conducted locally following previously published pipelines [75]. First, individual T1-weighted images were parcellated by transforming the Brainnetome (BN) atlas [76], which comprises 210 cortical and 36 subcortical regions, to native space through volumetric segmentation (FSL), yielding 246 regions-of-interest (ROIs). These ROIs were then used for both the structural and functional analyses. Probabilistic tractography was performed on diffusion MRI data using MRtrix3 to generate whole-brain tractograms [117]. The tractography procedure involved seeding ten million streamlines at the grey matter-white matter interface, with streamlines filtered according to length constraints and anatomically-constrained tractography (ACT) priors [118]. The resulting tractography streamlines were then combined with the parcellation scheme to construct 246 × 246 structural connectivity matrices where the connection strength between each pair of regions was quantified using the mean fractional anisotropy (FA) along their interconnecting white matter pathways. For functional time series extraction, the 246-region parcellation was resampled from each subject’s T1-weighted space to their preprocessed fMRI space using FreeSurfer. Regional time series were then obtained by averaging the BOLD signals across all voxels within each parcellated region, resulting in a 246 × 490 time series matrix (regions × timepoints). For subsequent analyses, we focused exclusively on cortical regions and used individual structural connectivity (210 cortical regions × 210 cortical regions) and functional time series (210 cortical regions × 490 timepoints).

### Extraction of brain states

We employed the method developed by Ceballos et al. [48] to identify seven distinct brain states from the resting-state fMRI data for each participant. This approach offers two significant advantages: it effectively captures individual-specific brain activation patterns while maintaining comparability and generalizability across studies due to its dataset-independent framework.

Specifically, using the Brainnetome atlas [76], we mapped 210 cortical regions to the seven intrinsic networks defined by Yeo et al [77]: visual (VIS), somatomotor (SOM), dorsal attention (DAT), ventral attention (VAT), limbic (LIM), frontoparietal (FPN), and default mode (DMN) networks. From the processed resting-state fMRI data, we extracted BOLD activity time series (490 time points) for all the cortical regions and performed z-score normalization across time for each region. Network-level activity was then derived by averaging signals from regions within each intrinsic network. To identify the dominant networks, we compared the network activity levels at each time point, designating the network with the highest activity as dominant. Time points where no network activity exceeded the mean activity level (zero in the z-scored data) were excluded to ensure robust state identification (results without this thresholding are provided in Figs. S3-6). Finally, representative brain states were generated by averaging the regional activity patterns across the time points of network dominance, capturing the characteristic magnitude and sign of cortical activity associated with each intrinsic network’s activation (Fig. 2a).

### Temporal brain state dynamics

After identifying seven discrete brain states, we characterized their temporal dynamics by computing four related metrics [18, 52] for each subject to investigate the differences between the MDD patients and the HCs. Fractional occupancy (FO) quantifies the temporal proportion of each state by calculating the ratio of the number of volumes assigned to a particular state to the total number of volumes. Dwell time (DT) captures the temporal persistence of each state by measuring the average duration (in seconds) of consecutive volumes within the same state. Appearance rate (AR) characterizes the frequency of state occurrence by computing the number of times a state was transitioned into per minute. Transition probability (TP) describes the patterns of state-to-state transitions by quantifying the likelihood of switching from one state to another.

### Control energy and energy regulation capacity

Here, we utilized network control theory (NCT) to model the energy required for specific brain state transitions with the constraints of the underlying structural connectome (approximated by structural connectivity), following prior work [10, 11, 21, 36, 42, 43, 50]. While this procedure has been detailed elsewhere [21, 43, 50], we summarize it briefly here.

With a representative *N* × *N* structural connectivity *A,* obtained as described above, where *N* is the number of regions, and a vector *x*(*t*) of size *N* × 1 represents the state of the system at time *t*, the temporal evolution of network activity can be modeled as a linear function of its connectivity,

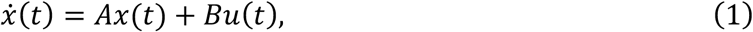

where *B* is an *N* × *N* matrix defining the control nodes and control input weights, and *u*(*t*) is the time-dependent control signal. Using optimal control, the controlled path from an initial state *x*_0_ to the final target state *x_T_* was designed to follow an optimal trajectory that balances the energy needed for the transition with the distance to the desired target state [21], i.e.,

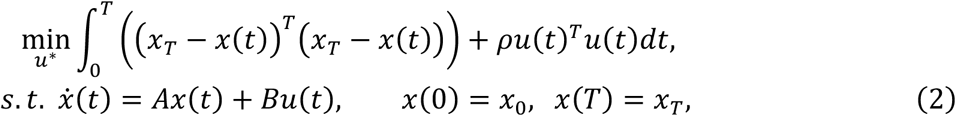

and the optimal control energy of a trajectory, integrated over time *T*, is defined as

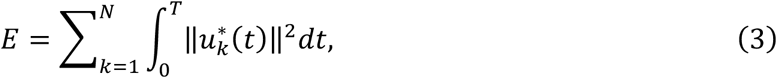

where *T* is the control horizon, and ρ is the penalizing parameter. In this context, the control energy required to remain in a given state is termed “persistence energy,” while “transition energy” refers to the control energy associated with transitions between different states. Following prior work, we set *T* = 1, and ρ = 1 (Results for T=3 are shown in Figs. S4-5).

From the control energy, we can also obtain the control stability, which is defined as the inverse persistence energy needed to maintain a state [50],

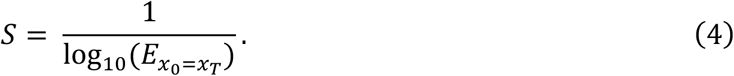

Based on it, we introduced a global stability, which is calculated as the inverse average persistence energy across different, considered states.

Building on NCT, we modeled two types of brain state control and estimated the associated energy costs, with control inputs taking the form of endogenous signals associated with internal cognitive demand or mental load [43, 49, 50]. The first of these probed the brain’s global energy efficiency with uniform control input weights, treating all brain regions equally in terms of their influence on system dynamics (Fig. 3). In this context, we denoted the average transition energy across different state pairs as *aveTE_0_*. The second one incorporated region-specific influences to quantify the capacity of a certain region in regulating energy with region-specific heterogeneous control input weights (Figs. 1 and 4). Specifically, we increased the control weight of a specified brain region *k* by one while keeping the control weights of all the other regions unchanged and then calculated the average transition energy across different state pairs as was done with uniform input weights, denoted as *aveTE_k_*. Consequently, the regional energy regulation capacity of region *k* was calculated as 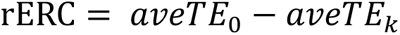. By repeating this process for each brain region of interest, we obtained a region-wise vector of energy regulation capacity (Fig. 1b).

### Association between regional energy regulation capacity and depressive symptoms

We conducted a partial least squares (PLS) correlation analysis [83] to examined the relationship between rERC patterns in MDD and the severity of depressive symptoms (quantified by PHQ-2 scores). First, we computed the correlation matrix between rERC values across brain regions and behavioral variables (PHQ-2 scores), with age, sex, and assessment center included as covariates. Singular value decomposition was then applied to this correlation matrix to identify the latent components. The significance of these components was assessed using permutation testing (10000 permutations), and the stability of the brain and behavior weights was evaluated through bootstrapping (5000 iterations). For the significant components, we examined the spatial correlations between regional loadings and the case-control *t*-map using spin tests to account for spatial autocorrelation. See Supplementary Results for details.

### Association analysis between energy regulation and neurotransmitter profiles

We examined the spatial association between the case-control rERC *t*-map and neurotransmitter distribution maps. The neurotransmitter receptor/transporter density maps were obtained from a comprehensive cortical profile derived from PET imaging studies of over 1,200 healthy participants [84]. From this, we extracted 19 distinct receptor and transporter maps spanning nine neurotransmitter systems: dopamine, norepinephrine, serotonin, acetylcholine, glutamate, GABA, histamine, cannabinoid, and opioid (see Supplementary Materials). These maps were parcellated to 210 cortical regions using the same atlas as the rERC *t*-map. Spearman correlations were calculated between the rERC *t*-map and each neurotransmitter receptor/transporter density map, with spin tests (see Methods: spin test) applied to correct spatial autocorrelation. Statistical significance was determined after Bonferroni correction for multiple comparisons.

### Association analysis between energy regulation and gene expressions

Brain-wide gene expression profiles were obtained from the Allen Human Brain Atlas (AHBA) [85, 86]. The microarray expression data were preprocessed using *abagen* [119] and mapped to the Brainnetome atlas [76]. After removing genes with low similarity across donors (*r* < 0.2), 5,826 genes were retained for subsequent analyses.

We employed PLS regression [68] to examine the spatial association between gene expression patterns and the rERC *t*-map. Regional gene expression levels served as predictor variables, with rERC *t* values as the response variable. The significance of the PLS components was assessed using permutation testing to test the null hypothesis that the PLS components explained no more covariance between the rERC *t* map and brain-wide gene expression than expected by chance [120]. Meanwhile, to account for spatial autocorrelation, of the rERC *t*-map (10000 random rotations), spherical rotations were considered (see Methods: spin test). Bootstrapping was used to estimate the normalized weights, Z scores, of each gene for each significant PLS component. Genes with |*Z*| > 3 were extracted for enrichment analysis using Metascape [87]. The significance threshold was set at *q* < .05, using the Benjamini-Hochberg correction to account for multiple testing. Detailed procedures are provided in the Supplementary Materials.

Moreover, following the methodology described in Hansen et al. [121], we performed a cell-type deconvolution analysis to examine cell-type enrichment in the gene sets identified by the PLS analysis. We utilized the cell-specific gene sets derived from five human adult postmortem single-cell and single-nucleus RNA sequencing studies [122–125], which identified seven major cortical cell types: astrocytes, endothelial cells, microglia, excitatory neurons, inhibitory neurons, oligodendrocytes, and oligodendrocyte precursors [126]. For each cell type, we calculated the ratio of genes that were preferentially expressed in that cell type. The statistical significance was evaluated against null distributions generated by repeating 10,000 times using randomly selected gene sets of equal size with two-tailed tests and Bonferroni correction for multiple comparisons.

### Spin tests

To account for spatial autocorrelation in cortical measures, we employed a permutation-based null model, termed a spin test [127, 128]. This approach involves projecting the parcel coordinates onto a spherical surface and applying random rotations to generate null distributions. Specifically, each cortical region was assigned the value of its nearest neighbor after rotation, with this process repeated many times [129]. This method preserves both the distribution of cortical values and the inherent spatial relationships in the data.

### Statistical Analysis

Group differences between the MDD and HC participants were assessed using two-sample *t* tests for temporal characteristics (fractional occupancy, dwell time, appearance rate, and transition probability), energy metrics (transition energy, persistence energy, average transition energy, and stability), and regional energy regulation capacity, with sex, age, and assessment center included as covariates. All group comparison results were corrected for multiple comparisons using the Benjamini-Hochberg correction (*p*_FDR_ <.05). Regional correlations were evaluated using Spearman’s correlation coefficient to ensure robustness against potential outliers. Spatial autocorrelation was addressed using spin tests as described above.

## Data Availability

The data supporting the findings of this study are available from the UK Biobank. Access to these data is subject to restrictions, as they were utilized under a specific license for this study. Data can be accessed at https://www.ukbiobank.ac.uk/ with the permission of UK Biobank. Human gene expression data that support the findings of this study are available in the Allen Brain Atlas (“Complete normalized microarray datasets”, https://human.brainmap.org/static/download). The cell type–specific genes are listed at https://github.com/jms290/PolySyn_MSNs/blob/master/Data/AHBA/celltypes_PSP.csv. The Brainnetome atlas can be downloaded at https://atlas.brainnetome.org/download.html. PET receptor and transporter maps are available at https://github.com/netneurolab/hansen_receptors. Source data supporting the findings of this study are available with reasonable request to the corresponding author.

## Code Availability

The processing pipeline for structural connectome construction and functional time series extraction was adapted from publicly available code (https://github.com/sina-mansour/UKB-connectomics). The code for extracting brain states can be found at https://github.com/NeuroenergeticsLab/control_costs. The code for computing the temporal characteristics of the brain states can be found at https://github.com/singlesp/energy_landscape. Control energy calculation is freely available at https://github.com/LindenParkesLab/nctpy. The code for performing two sample *t* tests with covariates can be found at https://github.com/Chaogan-Yan/DPABI/tree/master/StatisticalAnalysis. The partial least squares correlation analysis was implemented using a modified version of publicly available code at https://github.com/danizoeller/myPLS by including spin-based permutation tests for addressing spatial autocorrelation. The partial least squares regression for gene analysis is available at https://github.com/KirstieJane/NSPN_WhitakerVertes_PNAS2016. The code for spherical rotation and spin-based permutation testing can be found at https://github.com/frantisekvasa/rotate_parcellation and https://github.com/MICA-MNI/ENIGMA/tree/master/enigmatoolbox/permutation_testing. Gene enrichments were analyzed at https://metascape.org/gp/index.html#/main/step1. The code for cell type deconvolution is available at https://github.com/netneurolab/hansen_genescognition/blob/master/CTD/fcn_ctd.m. The Brain Connectivity Toolbox used to generate degree-preserving null models is freely available at https://sites.google.com/site/bctnet. The code used to generate the geometry-preserving null networks is freely available at https://www.brainnetworkslab.com/coderesources. The Neuromaps toolbox (version 0.0.5) for fetching brain maps is freely available at https://github.com/netneurolab/neuromaps. The ENIGMA Toolbox (v2.0.1) is freely available at https://github.com/MICA-MNI/ENIGMA. Custom code will be available with reasonable request upon publication.

## Acknowledgments

The research was conducted using the UK Biobank resources, with approved project number 88660. We sincerely appreciate the participants for their contributions and the UK Biobank team for their dedication to data collection, processing, and dissemination. We are grateful to our colleagues at the Brainnetome Center for their support and insightful discussions, as well as R. E. Perozzi and E. F. Perozzi for their valuable assistance in reviewing and refining the English and content of this paper.

This work was supported by the STI2030-Major Projects (Grant No. 2021ZD0200200), National Natural Science Foundation of China (Grant No. 12301642 to H.X., No. 62403465 to S.W.) and China Postdoctoral Science Foundation (GZC20232999 and 2024M753502).

## Author contributions

Q.L. and H.X. conceptualized the study. Q.L., X.H., and W.S. developed the methodology. Q.L., W.S., S.D. and X.L. curated the data. Q.L., H.X., W.S., S.D., X.C., N. Liu, N.L., and Y.Z. conducted the investigation. Q.L. and W.S. contributed to the visualization. T.J. supervised the study. Q.L. and H.X. drafted the original manuscript. Q.L., X.H., W.S., and T.J. reviewed and edited the manuscript.

